# Twist regulates Yorkie to guide lineage reprogramming of syncytial alary muscles

**DOI:** 10.1101/2020.06.22.165506

**Authors:** Marcel Rose, Jakob Bartle-Schultheis, Katrin Domsch, Ingolf Reim, Christoph Schaub

## Abstract

The genesis of syncytial muscles is typically considered as a paradigm for an irreversible developmental process. Notably, transdifferentiation of syncytial muscles is naturally occurring during *Drosophila* development. The ventral longitudinal heart-associated musculature (VLM) arises by a unique mechanism that revokes the differentiated fate from the so-called alary muscles and comprises at least two distinct steps: syncytial muscle cell fragmentation into single myoblasts and direct reprogramming into founder cells of the VLM lineage. Here we provide evidence that the mesodermal master regulator *twist* plays a key role during this reprogramming process. Acting downstream of *Drosophila* Tbx1 (Org-1) in the alary muscle lineage, Twist is crucially required for the derepression of the Hippo pathway effector Yki and thus for the initiation of syncytial muscle dedifferentiation and fragmentation. Subsequently, cell-autonomous FGFR-Ras-MAPK signaling in the resulting mono-nucleated myoblasts is maintaining Twist expression, thereby stabilizing nuclear Yki activity and inducing their lineage switch into the founder cells of the VLM.

## Introduction

Lineage restriction is an essential concept in multicellular organisms. On the one hand, a lower fate commitment enables a high regenerative potential, but on the other hand, a higher fate commitment prevents uncontrolled growth and cancer. During *Drosophila* embryonic development early fate decisions within the blastoderm are initiated by the bHLH transcription factor Twist, which is crucial for mesoderm specification (Thisse et al., 1988) and is required for the activation of mesoderm specific factors (Furlong et al., 2001; Leptin, 1991) such as the Drosophila FGFR *heartless* (*htl*), required for mesodermal cell migration (Beiman et al., 1996; Gisselbrecht et al., 1996; Shishido et al., 1993; Shishido et al., 1997), and *Mef2* initiating muscle differentiation (Bour et al., 1995; Lilly et al., 1995; Taylor et al., 1995). Although Twist was reported to promote embryonic myogenesis in *Drosophila* it appears to play the opposite role during vertebrate myogenesis, where primary Twist function is proposed in preventing the premature differentiation of myogenic cells (Baylies and Bate, 1996; Hebrok et al., 1994; Rohwedel et al., 1995; Spicer et al., 1996).

*Drosophila twist* is known to become restricted after its early mesodermal expression to adult muscle precursors (AMPs), muscle stem cells from which the adult somatic musculature will arise during metamorphosis (Bate et al., 1991). Herein we report that *twist* expression additionally persists in the embryonic and larval heart-associated syncytial alary muscles (AMs), which connect the embryonic heart to internal organs and the exoskeleton (Bataillé et al., 2020; Boukhatmi et al., 2014). Our recent work has shown that during metamorphosis a subset of these AMs exhibits an extraordinary level of cell fate plasticity. Escaping histolysis, the three anterior muscle pairs undergo an *in vivo* direct lineage reprogramming process that includes their dedifferentiation, fragmentation into mononucleated alary muscle derived cells (AMDCs) and re-differentiation into the ventral longitudinal musculature of the heart (VLM) (Schaub et al., 2015). We demonstrate that after the onset of metamorphosis *twist* expression persists during AM to VLM transdifferentiation in the AM lineage and that the reprogramming process crucially depends on Twist function. Mechanistically, the induction of trans-differentiation depends on the function of the *Drosophila* homolog of Tbx1 (*optomotor-blind-related-gene-1, org-1*), whereas the initiation of syncytial muscle dedifferentiation and fragmentation requires derepression of the transcriptional effector of the Hippo pathway, Yorkie (Yki) (Schaub et al., 2015; Schaub et al., 2019). We find that *twist* expression during lineage reprogramming is maintained by Org-1 and that Twist function is required for the derepression of Yki function in the anterior AMs. This identifies Twist as a crucial factor that links these two mechanisms. We show that Twist together with the *Drosophila* FGFR Heartless is essential for the reprogramming process of the AMDCs into the founder myoblasts of the VLMs and demonstrate that this lineage switch requires the stabilization of nuclear Yki localization in the AMDCs by active Ras-MAPK signaling. Altogether, our results show that a Twist/Yki/FGFR axis plays a central role in the regulatory network that mediates syncytial muscle cell fate plasticity and initiates a lineage switch during cellular reprogramming. Of note, vertebrate orthologs of Twist, Yki and FGFR are well known to be involved in the reprogramming processes that transform myogenic cells into rhabdomyosarcoma (Goldstein et al., 2007; Li et al., 2019; Tremblay et al., 2014). Our results connect the function of this set of regulators within a naturally occurring muscle lineage reprogramming event. Apart from adding mechanistic insights into this process they are potentially also relevant for neoplastic reprogramming.

## Results

### Twist is expressed in the alary muscle lineage and is required for AM to VLM lineage reprogramming

*Twist* expression is found during early embryonic development in the whole mesoderm, where its expression is regulated by upstream enhancer elements that are bound by the morphogen Dorsal (Jiang et al., 1991; Thisse et al., 1991). During later developmental stages somatic *twist* expression is known to be restricted to the AMPs that are specified via the rhomboid-triggered EGF signaling pathway (Bate et al., 1991; Figeac et al., 2010; Thisse et al., 1988). Unexpectedly, during an enhancer dissection analysis at the *twist* locus, reporter constructs driven by the complete downstream intergenic region (*twist*.*Ko6-GFP*, Fig. 1A) were found to include expression in the embryonic alary muscles (Dominik Müller, Jakob Bartle-Schultheis and Manfred Frasch, unpublished data; Fig. 1B). Visualization of the Twist protein together with an alary muscle-specific reporter (*tupAME-GFP*) during embryonic stages clearly show a localization of Twist in the nuclei of the embryonic alary muscles (Fig. 1C,C’). Moreover, *twist*.*Ko6-GFP* can be detected in larval alary muscles as well as in the VLM and the adult alary muscles (Fig. 1D-F). Live imaging demonstrated that the *twist*.*Ko6-GFP* reporter is active during the whole AM to VLM transdifferentiation process in the alary muscle lineage (video S1). The analysis of the expression pattern of a GFP-tagged version of Twist under endogenous control (*twi::GFP*) revealed that it is expressed in the *org-1-RFP* positive AMs, after dedifferentiation and fragmentation in the AMDCs as well as in the primordia of the VLM (Fig. 2A-C), indicating a potential role of Twist during alary muscle reprogramming. To dissect a possible requirement of *twist* function during VLM generation we induced its AM specific knock-down by RNAi or by inducible CRISPR using *org-1-GAL4* during metamorphosis. The analyses of the resulting phenotypes revealed a strong perturbation of VLM formation and alary muscle reprogramming (Fig. 2E,H). Accordingly, live imaging demonstrated that upon RNAi mediated *twist* knock-down alary muscle dedifferentiation and fragmentation is severely impaired (video S2). Past work has shown that Twist function during embryonic myogenesis in *Drosophila* as well as vertebrates depends on dimer choice and protein-protein interactions (Franco et al., 2011). Interestingly, the downregulation of the *Drosophila* E-protein Daughterless, which has been proposed to inhibit somatic myogenesis via heterodimerization with Twist (Castanon et al., 2001), by RNAi does not cause a significant phenotype during AM lineage reprogramming (Fig. 2H). In contrast, the elevation of the transcript levels of *twist* and *nautilus*, the *Drosophila* MyoD homolog (Michelson et al., 1990), using *org-1-GAL4* during lineage reprogramming abolishes VLM formation (Fig. 2F,G,H), indicating a dose-dependent, mechanistical role for bHLH factor interaction during transdifferentiation. These results demonstrate an important role for Twist function in mediating lineage plasticity and reversal of differentiated, syncytial muscle fate during AM lineage programming.

**Figure 1:**
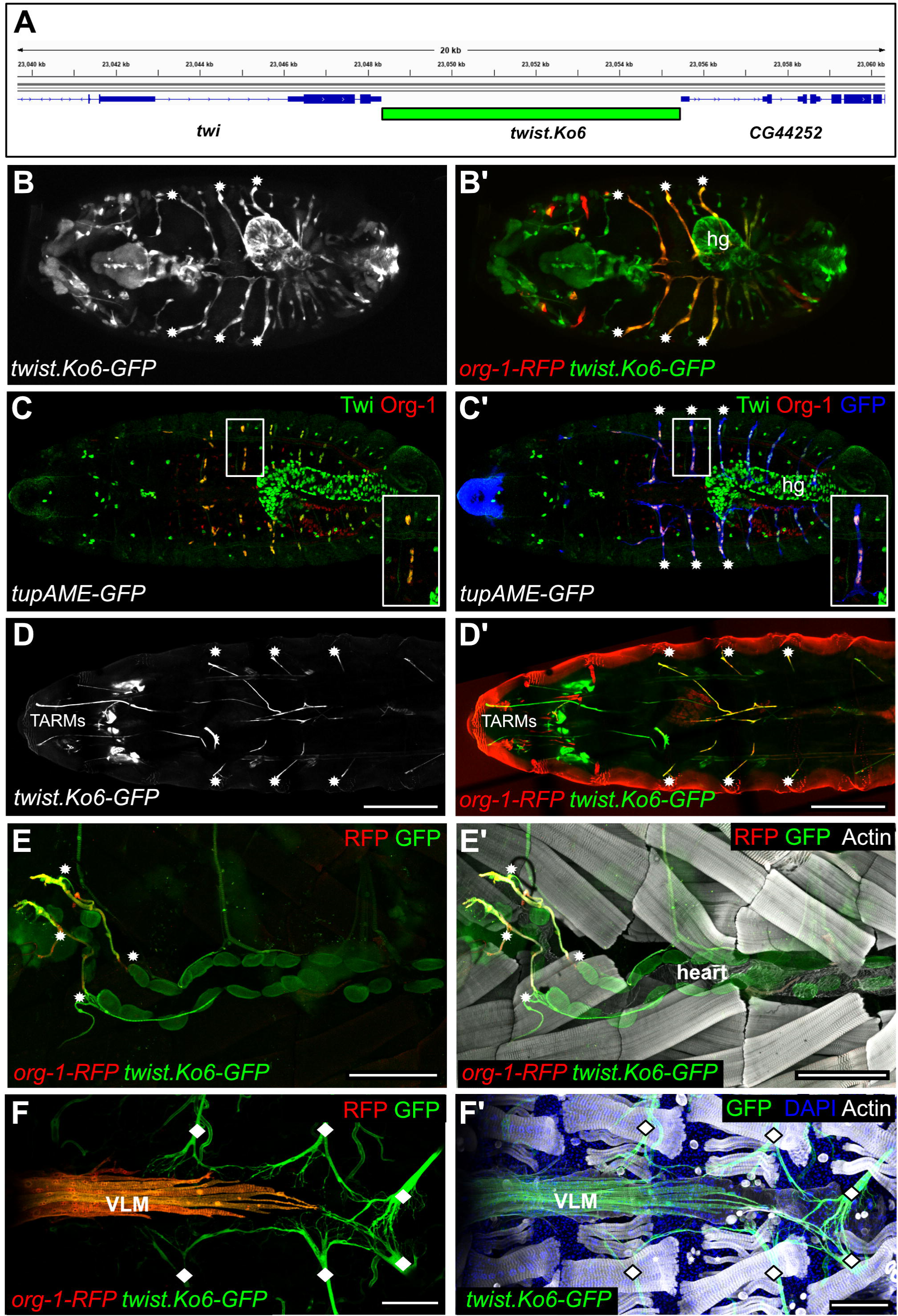
The bHLH transcription factor Twist is expressed in the syncytial alary muscles. **(A)** Schematic depiction of the genomic region used to create *twist*.*Ko6-GFP*. **(B-B’)** *In vivo* live imaging of an embryo carrying *twist*.*Ko6-GFP* and *org-1-HN18-RFP* (*org-1-RFP*). Strong GFP expression can be detected in the first three pairs of *org-1-RFP* positive alary muscles (asterisks). **(C-C’)** Dorsal view of an embryo carrying the alary muscle specific *tupAME-GFP* reporter and stained for Twist, Org-1 and GFP proteins. **(C)** Twist and Org-1 are clearly co-expressed in nuclei of the **(C’)** anterior three pairs of alary muscles (asterisks) as well as the posterior ones marked by the expression of *tupAME-GFP*. The magnified views show a single anterior alary muscle. **(D-D’)** *In vivo* live imaging of a dorsal view of a L3 larva carrying *twist*.*Ko6-GFP* and *org-1-HN18-RFP*. **(D)** *twist*.*Ko6* drives GFP expression in the thoracic alary related muscles (TARMs) and the first three pairs of **(D’)** *org-1-RFP* positive alary muscles. **(E-E’)** Visualization of the expression pattern of *org-1-RFP* and *twist*.*Ko6-GFP* in a dissected late L3 stage. RFP and GFP are colocalized in the anterior alary muscles (asterisks). **(F**,**F’)** *twist*.*Ko6-GFP* and *org-1-RFP* drive reporter expression in the ventral longitudinal musculature (VLM) attached to the adult heart of a stage P13 pharate fly. *Twist-Ko6-GFP* expression is additionally detected in the adult alary muscles (diamonds). Scale bars in D: 500 µm, Scale bars in E-F: 100 µm. Actin is visualized with phalloidin. DNA visualized with DAPI.

**Figure 2:**
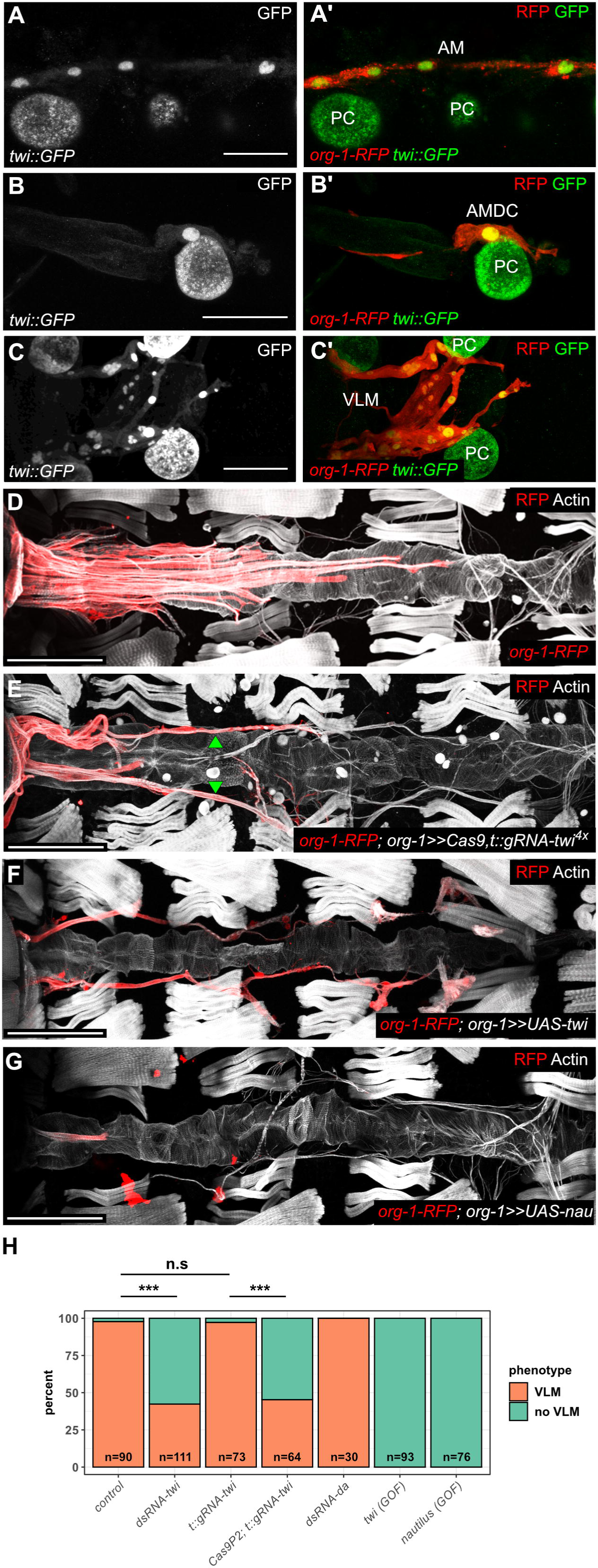
Twist is required for the lineage reprogramming of alary muscles into the VLM. **(A-C)** Visualization of the *org-1-RFP* lineage marker and an endogenously GFP tagged version of *twist* (*twi::GFP*). *twi::GFP* can be detected **(A-A’)** at the onset of metamorphosis in the nuclei of the syncytial alary muscles (AM) and **(B-B’)** during reprogramming in the nuclei of the formed mononuclear cells (AMDCs) as well as **(C-C’)** in the cells of the differentiating VLM. **(D)** *org-1-RFP* drives reporter expression in the ventral longitudinal musculature. **(E)** Induction of CRISPR in the alary muscles with *HN39-org-1-GAL4* (*org-1-GAL4*) against *twist* (*org-1*>>*Cas9; t::gRNA-twi*^*4x*^) blocks VLM formation and prevents AM fragmentation (arrowheads). **(F-G)** *HN39-org-1-GAL4* (*org-1-GAL4*) induced elevation of transcript levels of **(F)** *twist* (*org-1*>>*twi*) and **(G)** *nautilus* (*org-1*>>*nau*) leads to the abolishment of VLM formation. **(H)** Frequencies of observed VLM differentiation in the different genetic backgrounds. *org-1-GAL4-induced twist* RNAi (*dsRNA-twi*, n=111, p≤0.001) or CRISPR (*Cas9P2; t::gRNA-twi*, n=64, p≤0.001) abolishes significantly alary muscle transdifferentiation and VLM generation. Scale bars in A-C: 25 µm. Scale bars in D-E: 100 µm. Actin is visualized with phalloidin. DNA visualized with DAPI.

### Twist links Org-1 and Yki function during AM dedifferentiation

The induction of AM to VLM transdifferentiation crucially requires a transcriptional program that is activated by the AM lineage-specific induction of the *Drosophila* Tbx1 homolog Org-1 and its direct target, the Islet1 homolog Tailup (Schaub et al., 2015). To test for any functional connections between Org-1, Tup and Twist we analyzed the expression pattern of *twi::GFP* in genetic backgrounds that produce AM-specific *org-1* or *tup* knock-downs using *org-1-GAL4*. Strikingly, loss of Org-1 or of Tup leads to the nearly complete abolishment of *twi::GFP* expression in the *org-1-RFP* positive alary muscles (Fig. 3B,C), indicating that *twist* represents a downstream target of the Org-1 regulatory cascade during AM transdifferentiation. To further dissect the regulatory function of Org-1 on the *twist* cis-regulatory region (CRM) we reanalyzed the *twist*.*Ko6* downstream intergenic region and were able to identify an enhancer region that drives GFP reporter expression in the VLM lineage (*twiVLM-GFP*, Fig. 3D,F), suggesting a role of this CRM in *twist* regulation during AM lineage reprogramming. Org-1 as well as Tup are capable to significantly bind *twiVLM in vivo* (Fig. 3D) (Kudron et al., 2018) and the mutation of its predicted Org-1 binding sites (*twiVLM-orgI-IIImut-GFP*, Fig. 3E) leads to nearly complete abolishment of reporter activity in the VLM lineage (Fig 3G), thus demonstrating direct regulation of *twist* via this CRM by Org-1 during AM to VLM transdifferentiation.

**Figure 3:**
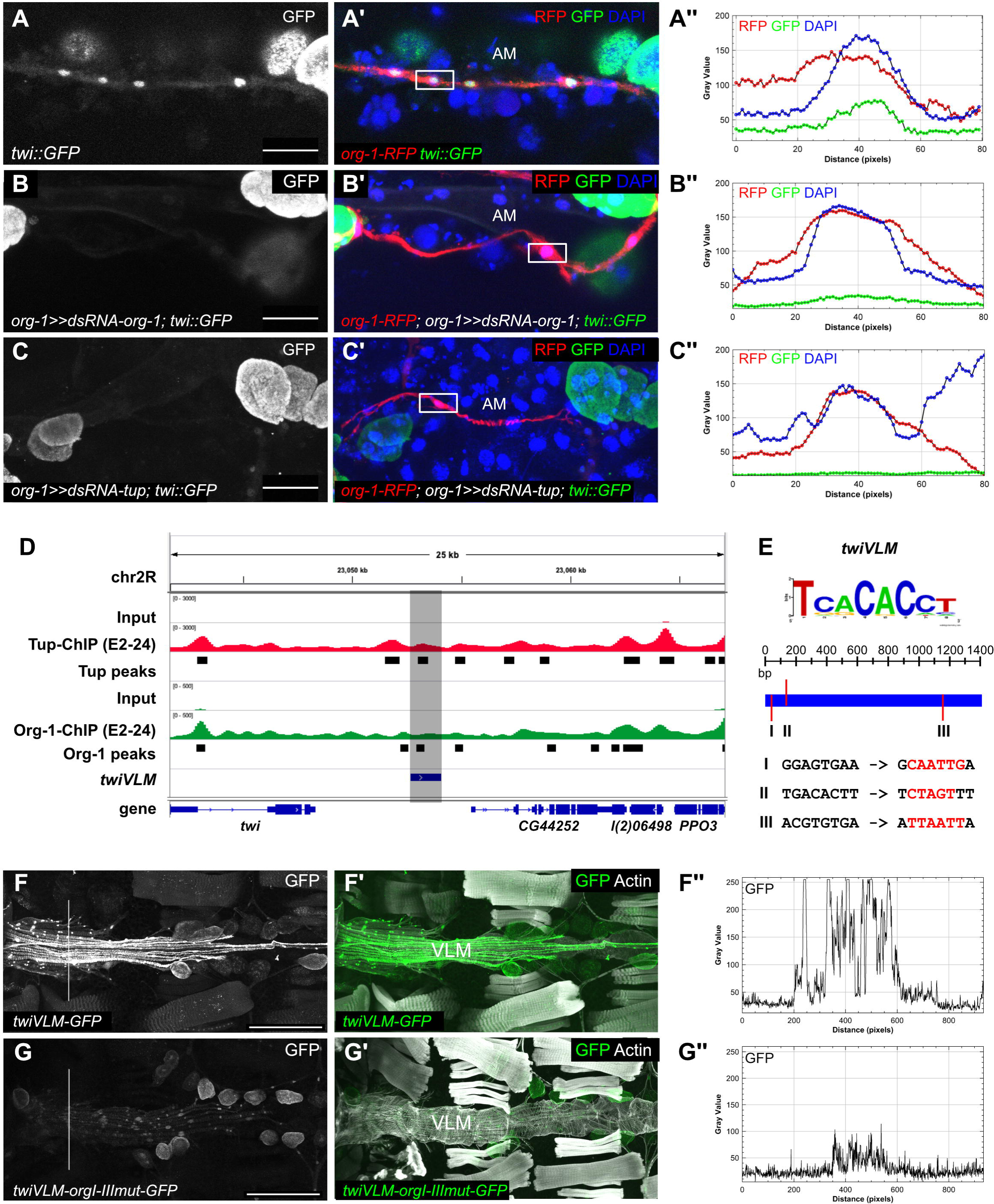
Twist expression during lineage reprogramming depends on Org-1 and Tup. **(A-A’’)** *twi::GFP* can be detected in the nuclei of the *org-1-RFP* positive alary muscles (AM) **(B**,**C)** Induction of RNAi with *HN39*-*org-1-GAL4* (*org-1-GAL4*) against **(B-B’’)** Org-1 (*org-1*>>*dsRNA-org-1*) or **(C-C’’)** Tailup (Tup) (*org-1*>>*dsRNA-tup*) leads to abolishment of *twi::GFP* expression in the alary muscles (AM). The indicated regions were used to quantify the expression level of nuclear DAPI, the *org-1-RFP* lineage marker and *twi::GFP* in **(A’’**,**B’’**,**C’’)** in the respective genetic backgrounds. **(D)** Illustration of the *twist* downstream genomic region showing Tup (red) and Org-1 (green) binding as well as called peaks during embryonic development (Kudron et al., 2018). The region used for the generation of *twiVLM-GFP* is indicated. **(E)** Schematic depiction of the *twiVLM* enhancer region and of the Org-1 binding motif. The predicted Org-1 binding sites and the point mutations that have been introduced to create *twiVLM-orgI-IIImut* are indicated. **(F-F’’)** *twiVLM-GFP* drives reporter expression in the VLM. **(G-G’’)**. The mutation of the predicted Org-1 binding sites in *twiVLM* (*twiVLM-orgI-III-GFP*) leads to the downregulation of GFP reporter expression in the VLM. The regions used for the quantification of GFP expression in **(F’’**,**G’’)** are indicated. Actin is visualized with phalloidin. DNA visualized with DAPI. Scale bars in A-C: 25 µm. Scale bars in F-G: 100 µm.

Yki represents the transcriptional effector of the Hippo network and its function as a transcriptional co-activator of Scalloped (Sd) is regulated by phosphorylation, thereby inhibiting its nuclear translocation (Goulev et al., 2008; Huang et al., 2005; Wu et al., 2008; Zhao et al., 2008). During AM to VLM transdifferentiation, Org-1 and Tup induce the derepression of the Yki/Sd complex in the syncytial AMs and thereby the activation of its target genes, thus leading to the dedifferentiation of the AMs into mononucleated cells and the induction of lineage reprogramming (Schaub et al., 2019). To dissect a functional connection between Twist and Yki during AM transdifferentiation we forced the co-expression of a phosphorylation resistant, constitutively active version of Yki together with RNAi against *twist* by using *org-1-GAL4*. Strikingly, we observed that constitutively active Yki caused a significant rescue of VLM formation in Twist knock-down backgrounds (*UAS-dsRNA-twi, UAS-yki*.*S168A;* p≤0.01; Fig. 4A-C), strongly suggesting that Twist is required for Yki derepression during alary muscle reprogramming. Taken together, these results support a mechanism that requires Org-1-dependent Twist function for the derepression of Yki in the alary muscles and concomitant activation of Yki/Sd target gene programs, which trigger alary muscle dedifferentiation.

**Figure 4:**
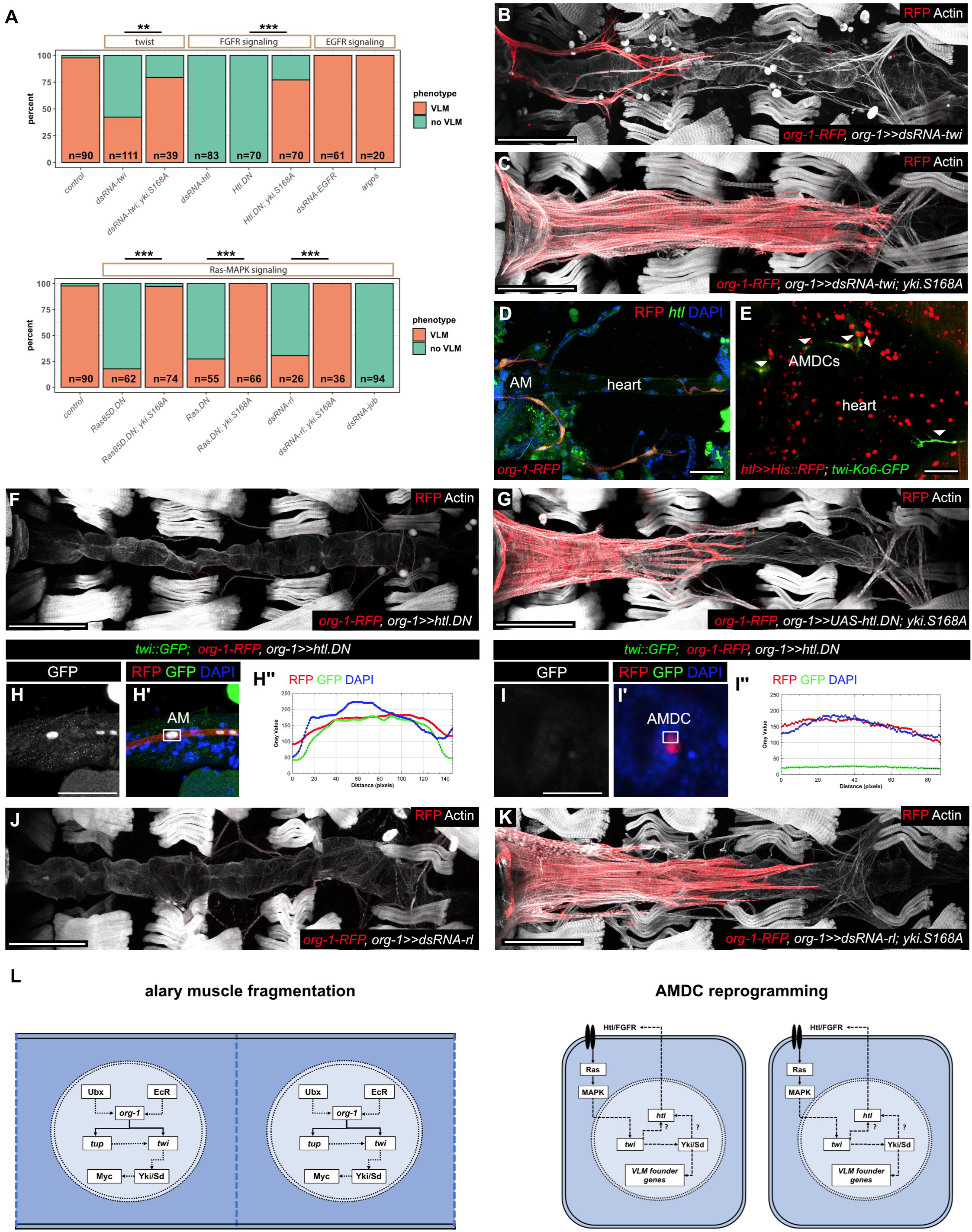
A Twist/Yki/FGFR regulatory axis mediates the induction of alary muscle fragmentation and the lineage switch during AMDC reprogramming. **(A)** Frequencies of observed VLM differentiation in the different genetic backgrounds. Co-expression of Yki^S168A^ can significantly rescue the phenotypes provoked by downregulation of Twist (n=39, p≤0.01) as well as forced expression of Htl^DN^ (n=54, p≤0.001), Ras85D^DN^ (n=74, p≤0.001), mammalian Ras^DN^ (n=66, p≤0.001) and knockdown of MAPK/Rolled (Rl) (n=36, p≤0.001). **(B)** Induction of RNAi against *twist* with *org-1-GAL4* (*org-1*>>*dsRNA-twi*) disrupts VLM differentiation and AM transdifferentiation. **(C)** VLM generation can be rescued by co-expression of phosphorylation resistant hippo pathway effector Yki^S168A^ in *twist* loss-of-function background (*org-1*>>*dsRNA-twi; yki*.*S168A)*. **(D)** *heartless* (*htl*) transcript can be detected in the *org-1-RFP* positive alary muscles (AM) at the onset of transdifferentiation. **(E)** Live imaging of flies carrying *htl-GAL4* and *UAS-His::RFP* in addition to *twi*.*Ko6-*GFP. Arrowheads indicate *htl-GAL4*-driven expression of Histone-RFP in GFP positive AMDCs. **(F)** Forced expression of a dominant negative version of Htl (*org-1*>>*htl*.*DN*) abolishes VLM differentiation. **(G)** These phenotypes can be significantly rescued by co-expression of Yki^S168A^ in Htl^DN^ background (*org-1*>>*htl*.*DN; yki*.*S168A*). **(H-H’’)** *org-1-GAL4* mediated induction of a dominant negative version of Htl (*org-1*>>*htl*.*DN*) has no effect on nuclear localization of GFP tagged Twist *(twi::GFP*) in alary muscles (AM) but **(I-I’’)** leads to abolishment of the *twi::GFP* expression in the AMDCs. **(H’’**,**I’’)** The indicated regions were used to quantify the expression level of nuclear DAPI, the *org-1-RFP* lineage marker as well as *twi::GFP* in the respective genetic backgrounds. **(J)** Induction of RNAi against MAPK/Rl with *org-1-GAL4* (*org-1*>>*dsRNA-rl*) disrupts VLM differentiation. **(K)** The phenotypes provoked by Rl loss-of-function can be rescued by co-expression of phosphorylation resistant Yki^S168A^ (*org-1*>>*dsRNA-rl; yki*.*S168A)*. **(L)** Schematic depiction of regulatory interactions during AM to VLM transdifferentiation. Continuous lines indicate direct regulation, dashed lines indicate indirect regulation and question marks proposed regulatory interaction. Twist expression is induced by the Ultrabithorax (Ubx)/Ecdysone Receptor (EcR)/Org-1/Tup transcriptional framework, thus leading to the de-repression of Yki/Sd and resulting in the activation of targets (e. g. Myc) required for fragmentation of the alary muscles. In the alary muscle derived cells (AMDCs), the cell autonomous activation of FGFR-MAPK signaling leads to the stabilization of Twist expression. This leads to the sustained de-repression of Yki/Sd, inducing the reprogramming process and the lineage switch by the activation of the VLM founder cell program. Scale bars in B,C&F,G&J,K: 100 µm. Scale bars in D,E: 50 µm. Scale bars in H,I: 25 µm. Actin is visualized with phalloidin. DNA is visualized with DAPI.

### Nuclear Twist and Yki are stabilized by FGFR signaling during muscle lineage reprogramming

During the establishment of the adult abdominal somatic musculature the Twist positive adult muscle precursors show expression of the *Drosophila* FGFR Heartless (Htl), which mediates the induction of founder myoblast fate in some of them (Dutta et al., 2005). Stainings for *htl* transcript as well as live imaging of the activity of a GAL4 line (*htl-GAL4*) (Jenett et al., 2012) driven by an *htl* enhancer containing previously characterized early embryonic Twist-binding sites (Stathopoulos et al., 2004; Zinzen et al., 2009), revealed that *htl* is expressed during AM to VLM transdifferentiation in the alary muscles as well as in the AMDCs (Fig. 4D,E; Fig. S1). Since after induction of RNAi against *htl* (*UAS-dsRNA-htl*) or the forced expression of a dominant negative version of Htl (*UAS-htl*.*DN*) no *org-1-RFP* positive muscles remain in pharate adults (Fig. 4A,F; Schaub et al. 2015), we deduce that activated FGFR signaling induces molecular events that are crucially required for alary muscle lineage reprogramming. To further investigate the role of FGFR signaling during AM to VLM transdifferentiation we performed live imaging experiments, thus demonstrating that mis-expression of a dominant negative version of Htl (*UAS-htl*.*DN*) using *org-1-GAL4* has no influence on alary muscle dedifferentiation and fragmentation but inhibits reprogramming of the *org-1-RFP* positive AMDCs into the founder cells of the VLM (video S3). Since reprogramming of the AMDCs crucially depends on the activation of Yki targets (Schaub et al., 2019), we asked if Yki activity in the AMDCs may be connected with activated Htl signaling. Our analyses revealed that the loss of VLM phenotypes generated via the activation of Htl^DN^ (*UAS-htl*.*DN*) can be significantly rescued by co-expression of a constitutive active Yki (*UAS-htl*.*DN; UAS-yki*.*S168A*, p≤0.001, Fig. 4A,F,G), thus suggesting that activated FGFR signaling is required for the maintenance of nuclear Yki function during AMDC reprogramming. Interestingly, Htl^DN^ (*UAS-htl*.*DN*) also leads to the loss of *twi::GFP* expression in the AMDCs, but not in the alary muscles before fragmentation (Fig. 4H,I). Taken into account that constitutive active Yki is epistatic over Twist (Fig. 4A,C), this suggests a regulatory interaction between FGFR, Twist and Yki function. Since it has been shown that Hippo signaling in wing discs can be negatively regulated via activated EGFR-Ras-MAPK signaling through the Ajuba LIM protein Jub (Reddy and Irvine, 2013), we speculated that sustained derepression of Yki in the AMDCs may depend on an analogous mechanism mediated by FGFR-Ras-MAPK signaling. Whereas induction of RNAi against EGFR or forced expression of the EGFR antagonist Argos has no phenotypic consequences (Fig. 4A), o*rg-1-GAL4* induced expression of dominant negative versions of *Drosophila* Ras (*UAS-Ras85D*.*DN*) as well as mammalian Ras proteins (*UAS-Ras*.*DN*) or downregulation of the *Drosophila* MAPK Rolled (Rl, *UAS-dsRNA-rl*) lead to the loss of VLM formation and AMDC reprogramming (Fig 4A,J). Strikingly, these phenotypes can be significantly rescued by co-expression of phosphorylation resistant Yki during AMDC reprogramming (p≤0.001, Fig. 4A,K), pointing towards a FGFR-Ras-MAPK-dependent regulation of nuclear Yki. The downregulation of Ajuba (*UAS-dsRNA-jub*) using *org-1-GAL4* provokes the abolishment of VLM generation (Fig. 4A), but unexpectedly these phenotypes cannot be significantly rescued by the constitutive activation of Yki targets, suggesting additional roles of Jub during AMDC reprogramming. Thus, we propose a model in which FGFR-Ras-MAPK signaling is required for the stabilization of nuclear Twist and Yki during reprogramming of the AMDCs into the VLM founder cells.

## Discussion

Twist function during embryonic muscle development in *Drosophila* has been described to possess dominant myogenic properties (Baylies and Bate, 1996), whereas vertebrate Twist is excluded from the myotome during embryogenesis and has been implicated in inhibiting myogenesis (Te and Reggiani, 2002). Twist expression and function in terminally differentiated striated, syncytial muscles has not been described yet. Here we show that Twist expression persist in the syncytial striated alary muscles after terminal differentiation and during metamorphosis in *Drosophila*. The alary muscles are a special type of striated muscles that are clearly distinguished from the body wall muscles (Bataillé et al., 2020; Boukhatmi et al., 2014), a feature that may be functionally connected to their persistent *twist* expression. During metamorphosis, the alary muscles demonstrate an extraordinary amount of cellular fate plasticity. Whereas the posterior larval alary muscles are remodeled into the adult alary muscles, the anterior ones generate the heart-associated ventral longitudinal musculature (VLM) by transdifferentiation (Schaub et al., 2015). Twist function and cellular plasticity have been connected in the literature mainly due to its role as one of the transcriptional master regulators during the transition from epithelial to mesenchymal states (EMT), crucially required for embryonic morphogenesis and often linked to metastasis in various cancer models (Lu and Kang, 2019). In terms of myogenic cell fate plasticity it is known that prolonged Twist expression negatively regulates muscle cell differentiation during embryonic myogenesis as well as adult flight muscle development in *Drosophila* (Anant et al., 1998; Cripps and Olson, 1998; Domsch et al., 2020), whereas forced expression of vertebrate Twist proteins can even reverse myogenic differentiation *in vitro* by unwiring the myogenic regulatory network from its targets (Hjiantoniou et al., 2008; Li et al., 2019; Liu et al., 2017; Mastroyiannopoulos et al., 2013) and appear to play a role during the regeneration of craniofacial muscles (Zhao et al., 2020). Here we provide evidence that Twist mediates the reversal of syncytial cell fate uncovering an unknown function of Twist in *Drosophila* muscles during metamorphosis.

It has been shown that sustained activation of the Hippo effector YAP inhibits the terminal myogenic differentiation program and that its *Drosophila* counterpart, Yki, induces reversal of syncytial differentiation during AM lineage reprogramming (Schaub et al., 2019; Sun et al., 2017; Watt et al., 2010). Here we demonstrate that Twist function is required for the derepression of Yki during AM transdifferentiation, connecting the Twist regulatory network to the activation of Yki target genes required for myogenic fate plasticity. Numerous studies have provided evidence for roles of YAP and TAZ, the vertebrate homologs of Yki, in EMT transdifferentiation and proposed that the induction of EMT upon forced Twist expression in cultured epithelial cells is connected to the inactivation of the Hippo pathway (Pan, 2010; Wang et al., 2016). These two mechanisms could be linked via the activation of JNK signaling and its transcriptional effector AP-1, both shown to be required for Twist-induced EMT as well as for Yki target activation (Nam et al., 2015; Sahu et al., 2015; Schaub et al., 2019; Sun and Irvine, 2013). Thus we speculate that Twist-induced abrogation of Yki target gene repression may represent a common molecular mechanism mediating cell fate plasticity in mesodermal cells. Twist function and FGFR signaling have been shown to be closely connected during development of mesodermal derivatives in various species (Beiman et al., 1996; Bloch-Zupan et al., 2001; Gisselbrecht et al., 1996; Harfe et al., 1998; Johnson et al., 2000; Pfeifer et al., 2014; Shishido et al., 1993; Shishido et al., 1997). During the genesis of the *Drosophila* adult musculature, FGFR signaling is required in Twist positive myoblasts for the specification of the abdominal founder cells that will seed the respective myofibers (Dutta et al., 2004; Dutta et al., 2005). We show that induction of FGFR signaling is required in another population of Twist positive myoblasts, i. e. those derived from the fragmentation of Twist positive syncytial alary muscles (AMDCs) during metamorphosis, to induce their lineage switch to the founder cells of the ventral heart-associated musculature (VLM). Furthermore, our data link active FGFR/MAPK signaling with Twist-mediated sustained de-repression of Yki in these cells. This connects Twist function to the induction of a potential Twist/Yki/FGFR feed-forward pathway driving AMDC lineage reprogramming. This is reminiscent of a mechanism proposed to be required for survival in cancer cells (Hua et al., 2016; Rizvi et al., 2016).

Taken together we propose a model in which Twist acts as a key regulator of alary muscle lineage reprogramming by coupling spatially and temporally specific transcriptional inputs to the de-repression of Yki targets (Fig. 4L). At the onset of metamorphosis, Org-1/Tbx1 and Tup/Isl1 expression are induced in the anterior alary muscles by the combinatorial inputs of the Hox protein Ultrabithorax and the ligand bound receptor of Ecdysone (Schaub et al., 2015). This leads to the stabilization of Twist expression only in the AMs fated to undergo transdifferentiation. Subsequently, the integration of the Org-1, Tup and Twist induced transcriptional networks leads to the de-repression of Yki/Sd target genes (e. g. *Myc*) required for the induction of muscle dedifferentiation and alary muscle fragmentation (Schaub et al., 2019). Once the alary muscles have fragmented into single myoblasts, the subsequent activation of FGFR/MAPK signaling in the AMDCs stabilizes the expression of Twist-induced Yki/Sd target genes, possibly via a regulatory Twi/Yki/FGFR feed-forward loop, and mediates reprogramming of the AMDCs into the founder cells of the VLM.

## Materials and methods

### *Drosophila* stocks and culture conditions

Standard *Drosophila melanogaster* genetics were performed at 25 °C. The following stocks were used: [*org-1-GAL4, org-1-RFP; twi::GFP*], [*org-1-RFP*; *twist*.*Ko6-GFP*], *tupAME-GFP* (Boukhatmi et al., 2014), [*org-1-GAL4, org-1-RFP*], [*org-1-GAL4, org-1-RFP*; *hand-GFP*], [*org-1-GAL4, org-1-RFP; UAS-Cas9*.*P2*], [*org-1-GAL4, org-1-RFP; UAS-yki*.*S168A*] (Schaub et al., 2019), *UAS-dsRNA-org-1* (62953), *UAS-dsRNA-tup* (51763), *UAS-dsRNA-twi* (51164), *UAS-htl*.*DN-B40; UAS-htl*.*DN*.*-B61* (5366), *UAS-dsRNA-htl-2* (35024), *UAS-dsRNA-EGFR-1* (60012), *UAS-dsRNA-EGFR-2* (25781), *UAS-argos*.*M*; *UAS-argos*.*M* (5363), *UAS-Ras85D*.*N17 (4845), UAS-Ras*.*N17* (4846), *UAS-dsRNA-rl* (34855), *UAS-dsRNA-jub* (32923), *GMR93H07-htl-GAL4* (40669), *UAS-His::RFP* (derived from 56555) and *twi::GFP* (79615) were obtained from the Bloomington *Drosophila* Stock Center. *UAS-dsRNA-htl-1* (6692) *UAS-dsRNA-da-1* (51297), *UAS-dsRNA-da-2* (51300) and *UAS-dsRNA-da-3* (105259) were obtained from the Vienna *Drosophila* Resource Center (VDRC).

### Generation of transgenic *Drosophila* stocks

To create *UAS-t::gRNA-twi*^*4x*^ we utilized the system described in (Port and Bullock, 2016). Using 1 ng pCFD6 (Addgene, Cat. No: 73915) as template, we amplified PCR fragments with the respective primers containing the gRNA sequences (Table S1) and utilizing Q5 Hot Start High-Fidelity 2X Master Mix (New England Biolabs, Cat. No: M0494L). After gel-purification of the PCR fragments with the QIAquick Gel Extraction Kit (Qiagen, Cat. No: 28704), these were assembled with BbsI-HF (New England Biolabs, Cat. No: 3539S) digested pCFD6 backbone to generate *UAS-t::gRNA-twi*^*4x*^ using the NEBuilder® HiFi DNA Assembly Cloning Kit (New England Biolabs, Cat. No: R5520S), following the instructions provided by the manufacturers.

To create *twiVLM-GFP*, the respective genomic region (2R:23052698..23054095) was amplified from *yw* DNA with the primers *KpnI-twiVLM-fw* (GGGGT ACCCCCAGTAAGGCAAATTGCTCAG) and *XhoI-twiVLM-rev* (CCGCTCGAGCGGAACTTGCCTTGTCCTTCGTC) utilizing Q5 Hot Start High-Fidelity 2X Master Mix (New England Biolabs, Cat. No: M0494L). The resulting PCR product was gel purified with the QIAquick Gel Extraction Kit (Qiagen, Cat. No: 28704) and cloned into *Kpn*I/*Xho*I of *pH-Stinger-AttB* (Jin et al., 2013) using T4 DNA ligase (New England Biolabs, Cat. No: M2020L).

For the analogous creation of *twiVLM-orgI-IIImut-GFP* the Org-1 binding sites within the *twiVLM* sequence were predicted with Target Explorer (Sosinsky et al., 2003) using a positional weight matrix generated by SELEX with Org-1 (Jin et al., 2013) and mutated via site-directed mutagenesis.

Transgenes of the constructs were generated by utilizing the phiC31/attP/attB system (Bischof et al., 2007) and performed by a commercial embryo injection service (BestGene Inc, Chino Hills, CA 91709, USA) to insert *UAS-t::gRNA-twi*^*4x*^ at the *attP40* landing site or to insert *twiVLM-GFP* as well as *twiVLM-orgI-IIImut-GFP* into the *attP2* landing site. Potential transformants were selected by mini-white expression.

### Fluorescent antibody staining

For antibody stainings of embryos the standard fixation protocol (Wieschaus and Nüsslein-Volhard, 1986) was applied. For dissection of pupal stages with microsurgery scissors the pupal cases were first removed and prefixation with 3,7 % formaldehyde for 30 minutes was performed. Pharate adults were dissected without prefixation. The dissected animals were fixed in 3,7 % formaldehyde for 10 minutes and were washed three times for 30 minutes in PBT (PBS + 0.1 % Tween©20).

Fixed embryos and pupae were blocked in 10 % Bovine Serum Albumin (Serva, Cat. No: 11930.04) for 1 hour. Samples were incubated for overnight (embryos) or two days (pupae) at 4 °C with the respective primary antibodies: Rabbit polyclonal anti-GFP (Rockland, Cat. No: 600-401-215, 1:500), Goat polyclonal anti-GFP (GeneTex Cat. No: GTX26673, 1:1000), Rabbit polyclonal anti-RFP (Rockland, Cat. No: 600-401-379S, 1:500), Rat polyclonal anti-Tropomyosin (MAC141, abcam, Cat. No: ab50567, 1:200), Rat polyclonal anti-Org-1 ((Schaub et al., 2012), 1:100), Guinea pig polyclonal anti-Twist (a gift of Katrin Domsch given by Eileen Furlong, 1:200), Rabbit polyclonal anti-Twist (a gift of Manfred Frasch, 1:500). To visualize filamentous actin Phalloidin conjugated either with Atto 488 (1:1000, Sigma-Aldrich, Cat. No: 49409) or Atto 647N (1:1000, Sigma-Aldrich, Cat. No: 65906) was added to the antibody solution. After the incubation with the primary antibodies the embryos and pupal dissections were washed three times for 30 minutes in PBT and incubated with the respective secondary antibodies: Alexa Fluor 488 conjugated Donkey Anti-Goat (abcam Cat. No: ab150129), Alexa Fluor 488 conjugated Goat Anti-Rabbit (ThermoFisher Scientific Cat. No: A-11008), DyLight 549 conjugated Goat anti-Rabbit (Jackson ImmunoResearch, Cat. No: 111-505-003) Alexa Fluor 555 conjugated Donkey Anti-Rabbit (abcam Cat. No: ab150074), Cy3 conjugated Donkey anti-Guinea pig (Jackson ImmunoResearch, Cat. No: 706-165-148), DyLight 649 conjugated Goat anti-Rat (Jackson ImmunoResearch, Cat. No: 112-495-003) in 1:200 dilution overnight at 4 °C. Finally, the stained animals were washed three times for 30 minutes in PBT and were embedded into Vectashield containing DAPI (Vectorlabs, Cat. No: H-1200).

### In situ hybridization chain reaction

To detect *heartless* mRNA in pupae we performed *in situ* hybridization chain reaction (HCR) as described in (Choi et al., 2018). For this purpose, we purchased a kit containing a custom *htl* DNA probe set and a fluorescently labelled DNA HCR amplifier as well as hybridization, wash and amplification buffers from Molecular Instruments (Los Angeles, CA 90041, USA).

For dissection of pupal stages with microsurgery scissors the pupal cases were first removed and prefixation with 3,7 % formaldehyde for 30 minutes was performed. The dissected animals were fixed in 3,7 % formaldehyde for 10 minutes, washed three times for 10 minutes in PBT (PBS + 0.1 % Tween©20), followed by a 5 minute wash in 50 % Methanol/PBT, a 5 minute wash in 100 % Methanol and another 5 minute wash in 50 % Methanol/PBT. The dissected pupae were rehydrated by three 5 minute washes in PBT followed by three 30 minute washes in PBT. After post-fixation with 3,7 % formaldehyde for 30 minutes the specimens were washed for three times with PBT for 5 minutes. After 30 minutes of prehybridization at 37°C in hybridization buffer, dissected pupae were hybridized with 0,4 pmol of the *htl* DNA probe set in 100 µl hybridization buffer for 16 hours at 37 °C. To remove unbound probes, the specimens were washed four times for 15 minutes with probe wash buffer at 37 °C, followed by three washes with 5xSSCT (5xSSC + 0.1 % Tween©20) at room temperature. For signal amplification the embryos and pupae were subjected to preamplification in amplification buffer for 30 minutes, followed by incubation with 6 pmol of each snap cooled, Alexa Fluor 647 labelled amplification hairpin in 100 µl amplification buffer for 16 hours at room temperature. To remove excess hairpins, two washes with 5xSSCT for 5 minutes and two washes with 5xSSCT for 30 minutes were performed. The pupae stained for *htl* mRNA localization were subjected to antibody stainings as described above.

### Sample analysis

Confocal Z-stacks of fixed or *in vivo* specimen were acquired with a Leica SP5 II (10x/0.4 HC PL APO Air, 20x/0.7 HC PL APO Glycerol, 63x/1.3 HC PL APO Glycerol). Maximum Projections of the Z-stacks were performed with FIJI/ImageJ (v1.52v) (Schindelin et al., 2012).

### *In vivo* time-lapse imaging

Stage P3 pupae of the respective genotypes were aligned on a strip of double-faced adhesive tape connected to a slide, covered with a drop of Halocarbon oil 700 (Halocarbon Products Corp, Sigma Aldrich Cat. No: H8898) and a coverslip. Time-lapse series were acquired on a Leica SP5 II confocal system using a HC PL APO 10x/0.4 Air. Acquisition was done over a time course of 48 hours with the following settings: Pinhole 1.4 AU, 1.2x optical zoom, scan speed 200 Hz, Z-stack about 30-35 sections with a step size of 2-4 µm and time intervals of 10 min per stack. Movies were generated using Leica Application Suite X (LAS X, Leica Microsystems) and with FIJI/ImageJ (v1.52v) (Schindelin et al., 2012).

### Statistical analysis

VLM phenotype frequency was quantified with living pharate stages of the respective genotypes that were aligned on a microscope slide. With the aid of a fluorescence equipped microscope (Nikon, Eclipse 80i) the *org-1-RFP* lineage marker in the living animals was visualized and phenotypes with fully differentiated VLM (proper aligned and elongated muscle fibers) were scored versus phenotypes ranging from missing VLM to defective VLM differentiation (no aligned muscle fibers). The Chi square test function of Microsoft Excel (Microsoft) was used for statistical analysis of the samples and statistical significance was defined by p≤0.05. Fluorescent signal quantification and illustration was performed with FIJI/ImageJ (v1.52v) (Schindelin et al., 2012).

## Acknowledgements

We thank the *Drosophila* community for providing us with antibodies and fly stocks and in particular Manfred Frasch (Friedrich-Alexander-Universität Erlangen-Nürnberg) for his gifts of unpublished materials. We are also grateful to the Bloomington *Drosophila* Stock Center and the Vienna *Drosophila* Resource Center for providing fly stocks as well as to the Optical Imaging Centre Erlangen (OICE) for their support and facilities.

## Author contributions

Conceptualization: Christoph Schaub; Investigation: Marcel Rose, Jakob Bartle-Schultheis, Katrin Domsch, Ingolf Reim and Christoph Schaub ; Writing – Original Draft: Christoph Schaub; Writing – Review & Editing: Marcel Rose, Jakob Bartle-Schultheis, Katrin Domsch, Ingolf Reim and Christoph Schaub; Funding Acquisition: Christoph Schaub; Supervision: Christoph Schaub

## Supplemental information

**Figure S1**.Activity of *htl* expression during AM/VLM transdifferentiation. Flies carrying *htl-GAL4* (GMR93H07) and *UAS-His::RFP* in addition to *twi*.*Ko6-*GFP were imaged during metamorphosis. **A-J** show stills from the same image series in developmental order (dorsal view). Arrows indicate *htl-GAL4*-driven expression of Histone-RFP in **(A, B)** either AMs prior to abdominal compaction or **(C-J)** in AMDCs and the developing VLM. (Note that only some portions are optically accessible from the outside particularly at early pre-pupal stages.) Expression is also observed in the central heart and various flanking abdominal skeletal muscles (both GFP-negative) as well as the developing flight muscles (GFP-positive, to the left in C-J).

**Video S1**.Dorsal view of a pupa that carries *twist*.*Ko6-GFP* and *org-1-RFP*. The movie starts at late stage P3 and demonstrates the co-expression of *twist*.*Ko6-GFP* and *org-1-RFP* during the transdifferentiation process of the larval alary muscles into the ventral longitudinal musculature of the heart. At the beginning of the movie, the syncytial *org-1-RFP* and *twist*.*Ko6-GFP* positive anterior alary muscles initiate dedifferentiation into mesenchymal, mononucleated myoblasts. These keep close contact to each other and migrate towards the dorsal vessel where they start re-differentiating into the ventral longitudinal heart muscles elongating along the heart tube towards the posterior. In addition, *twist*.*Ko6-GFP* can be detected in the remodeling *org-1-RFP* negative larval posterior alary muscles, the dorsal adult muscle precursors (AMPs) as well as the hindgut musculature.

**Video S2**.Dorsal view of a pupa that carries *hand-GFP* and *org-1-RFP* and in which *org-1-GAL4* mediated RNAi against *twist* was induced (*org-1*>>*UAS-dsRNA-twi*). The movie starts at late stage P3 and demonstrates that knock-down of *twi* prevents the alary muscles from dedifferentiating into mononucleated myoblasts, suggesting that Twist function is indispensable for the induction of this essential process during alary muscle trans-differentiation.

**Video S3**.Dorsal view of a pupa that carries *hand-GFP* and *org-1-RFP* and in which a dominant negative version of the *Drosophila* FGFR Htl was induced (*org-1*>>*UAS-htl*.*DN*). The movie starts at late stage P3 and shows that interference with FGFR mediated signaling has no impact on the fragmentation of the alary muscles into mononucleated myoblasts but demonstrates that transdifferentiation is arrested under these conditions after alary muscle dedifferentiation. This suggests an essential role of Htl-signaling during the lineage reprogramming of the alary muscle derived myoblasts into the founders of the VLM during transdifferentiation.

