## Supplementary figures and images for "Twist regulates Yorkie to guide lineage reprogramming of syncytial alary muscles"

### Figure S1

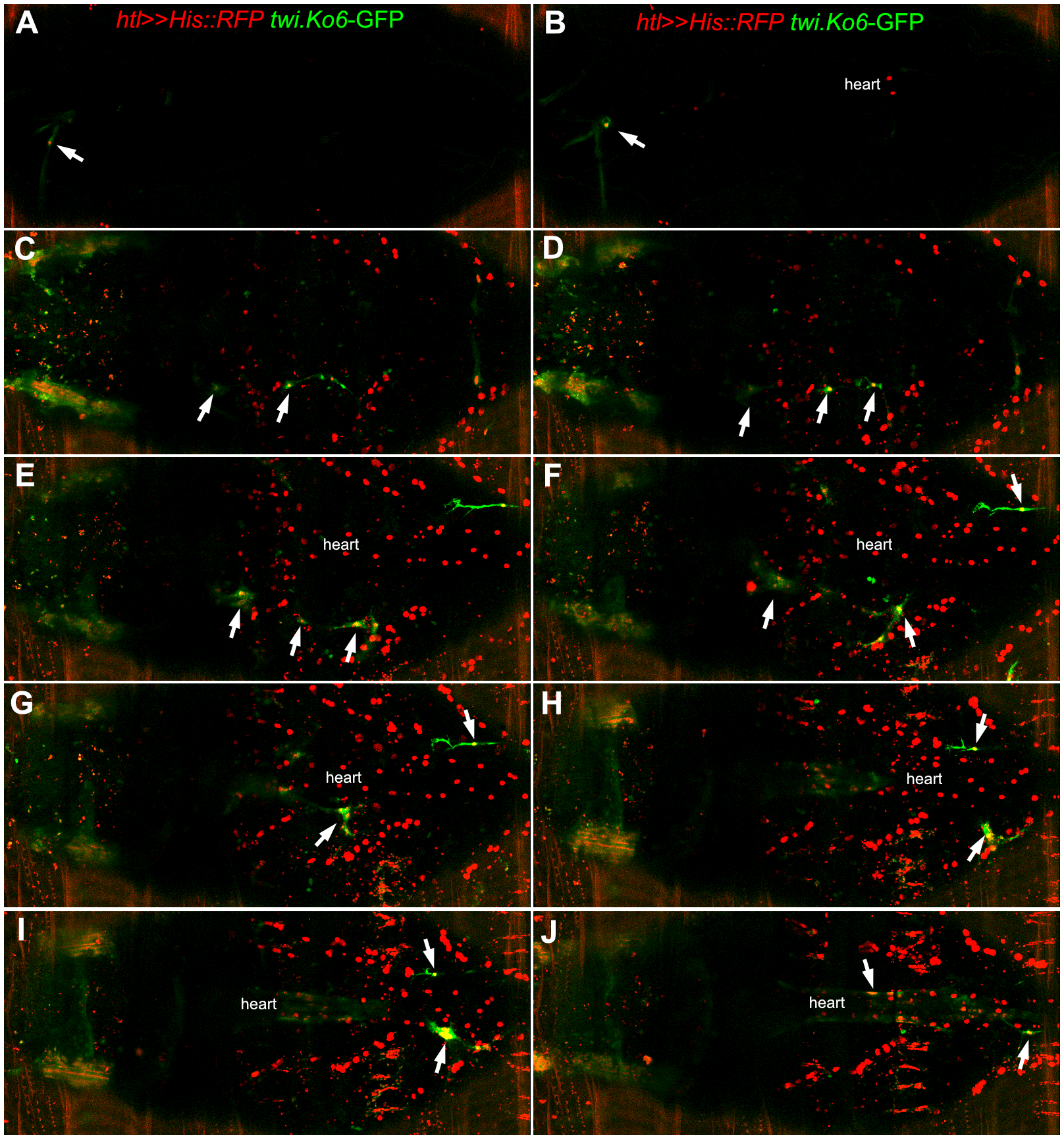
